# The *Arabidopsis* Diacylglycerol Kinase 4 is involved in nitric oxide-dependent pollen tube guidance and fertilization

**DOI:** 10.1101/665810

**Authors:** Aloysius Wong, Lara Donaldson, Maria Teresa Portes, Jörg Eppinger, José Feijó, Christoph Gehring

## Abstract

Nitric oxide (NO) is a key signaling molecule that regulates diverse biological processes in both animals and plants. In animals, NO regulates vascular wall tone, neurotransmission and immune response while in plants, NO is essential for development and responses to biotic and abiotic stresses [1–3]. Interestingly, NO is involved in the sexual reproduction of both animals and plants mediating physiological events related to the male gamete [2, 4]. In animals, NO stimulates sperm motility [4] and binding to the plasma membrane of oocytes [5] while in plants, NO mediates pollen-stigma interactions and pollen tube guidance [6, 7]. NO generation in pollen tubes (PTs) has been demonstrated [8] and intracellular responses to NO include cytosolic Ca^2+^ elevation, actin organization, vesicle trafficking and cell wall deposition [7, 9]. However, the NO-responsive proteins that mediate these responses are still elusive. Here we show that PTs of *Arabidopsis* lacking the pollen-specific Diacylglycerol Kinase 4 (DGK4) grow slower and become insensitive to NO-dependent growth inhibition and re-orientation responses. Recombinant DGK4 protein yields NO-responsive spectral and catalytic changes *in vitro* which are compatible with a role in NO perception and signaling in PTs. NO is a fast, diffusible gas and, based on our results, we hypothesize it could serve in long range signaling and/or rapid cell-cell communication functions mediated by DGK4 downstream signaling during the progamic phase of angiosperm reproduction.

## Results and Discussion

### *DGK4* is required for NO-dependent PT growth and re-orientation responses and affects reproductive fitness

Previously, the pollen-specific *Arabidopsis* Diacylglycerol Kinase 4 (DGK4) (TAIR ID: At5g57690) has been suggested to harbor a gas-sensing region [2] and DGK activity has been associated with important roles in pollen germination and growth [10]. Therefore, we chose to investigate the NO-dependent PT growth responses in wild-type (WT) (ecotype *Col-0*) plants and plants lacking DGK4. We characterized a homozygous *dgk4* plant with a T-DNA insertion 424 bp upstream of the *DGK4* gene (*dgk4-1*; SALK_151239) and observed a 50% reduction in *DGK4* expression (Figure S1). *dgk4-1* PTs grow significantly slower *in vitro* compared to WT across a pH range from 6.5 to 8.5 and in both optimal and reduced Ca^2+^ media (Figure 1A). After four hours of germination, WT PTs reached an average length of 176 *µ*m while *dgk4-1* only reached 144 *µ*m at pH 7.5 (*n* > 100; *P* < 0.05). This mutant line was also reported to have PTs with altered stiffness and adhesion properties [11]. When exposed to the NO donor sodium nitroprusside (SNP), both WT and the *dgk4-1* PTs show NO dose-dependent reduction of growth rates (Figure 1B) much like those reported in other systems like *Lilium longiflorum* [8] or *Camelia sinensis* [12]. But, importantly, PT growth rate of *dgk4-1* becomes insensitive to concentration increases over 50 nM SNP, while WT PT growth rate inhibition continues to decrease in a concentration dependent manner up to 200 nM SNP (Figures 1B and 1C). This differential sensitivity was also observed for another NO donor, DEA NONOate (Figure 1B). This result shows that PTs of *dgk4-1* are less responsive to NO thus suggesting a functional link between NO and DGK4. These phenotypes were confirmed in a second independent homozygous *dgk4* mutant plant (*dgk4-2*; SALK_145081, T-DNA insertion 268 bp upstream of the *DGK4* gene) (Figure S1). PTs of *dgk4-2* display a similar NO insensitivity in terms of growth rate when compared to WT (Figure S2). We further examined the effect of NO on directional growth of PTs using *dgk4-1*. When challenged with a NO point source (a SNP-loaded pipette tip), we observed that both WT and *dgk4-1* PTs showed a negative chemotropic response, bending away from the NO source much like the observations previously shown in lily [8]. However, PTs bending angles in WT (31.9 ± 2.4º; *n* = 4) are twice as sharp than those of *dgk4-1* (14.9 ± 2.5º; *n* = 4) (Figure 1D) revealing a desensitization in the perception of NO in the *dgk4-*1 mutant. Given that the NO critical concentration for this negative chemotropic reaction response was previously estimated to be in the range of 5-10 nM [8] this result is consistent with a signaling role for DGK4 in NO sensing.

**Figure 1.**
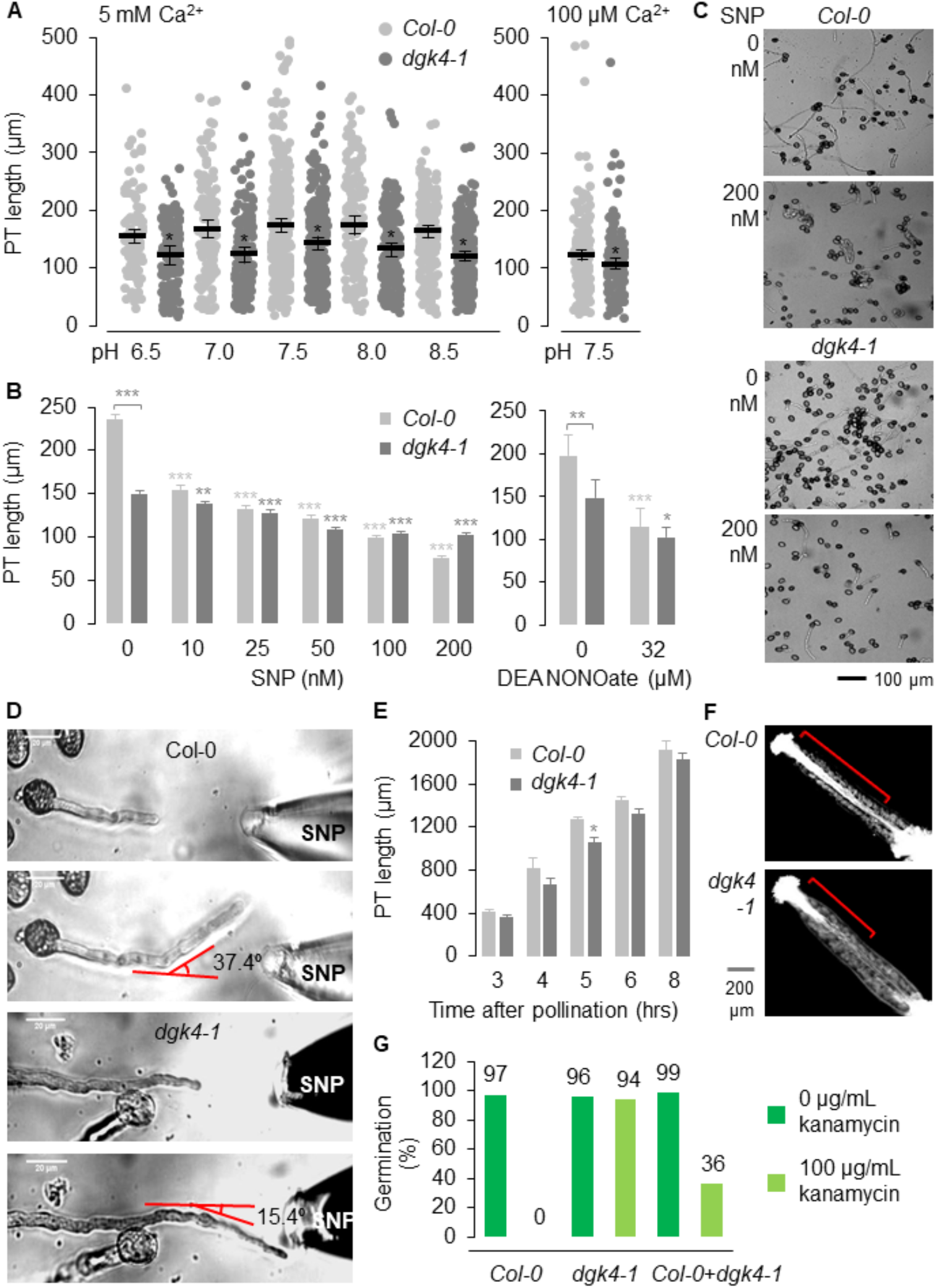
*dgk4-1* PT has reduced growth rate and NO-dependent PT growth responses. (**A**) The growth of *dgk4-1* PT is slower than that of *Col-0* consistent across a range of pH (6.5 to 8.5) and in both optimal (5 mM) (left) and reduced (100 µM) Ca^2+^ (right) media. Error bars represent standard error of the mean (*n* > 100). *\* = P* < 0.05 compared to PT length of *Col-0*. (**B**) NO-dependent inhibition of *dgk4-1* PT growth is reduced compared to that of *Col-0*. NO was provided by either SNP or DEA NONOate. (**C**) Representative images of *dgk4-1* and *Col-0* PTs with and without NO (provided by 200 nM SNP). *In vitro* pollen germination was performed as detailed previously [7, 8] and PT length was analyzed by capturing images covering the entire growth area of the culture dish that was mounted on an automated stage using the Nikon Eclipse TE2000-S inverted microscope equipped with a Hamamatsu Flash28s CMOS camera. PT lengths were then measured using NeuronJ [22]. Error bars represent standard error of the mean (*n* > 150). *\* = P* < 0.05*, ** = P* < 0.005 and *\*\*\* = P* < 0.0005 compared to PT length of untreated sample. (**D**) A representative image of the response of a growing *Col-0* PT bending away from a NO glass probe containing 1% agarose-SNP (10 mM) at a sharper angle than *dgk4-1*. The PT NO-dependent re-orientation response was monitored (*n* = 4) by real-time imaging using a Nikon Eclipse TE300 inverted microscope equipped with an Andor iXon3 camera and the bending angles measured using ImageJ [23]. (**E**) In the pistil, PT growth of *dgk4-1* is slowed. (**F**) The representative pistil image at 5 hours post-fertilization shows a higher density of longer *Col-0* PTs. Error bars represent standard error of the mean (*n* > 3). *\* = P* < 0.05 compared to tube length of *Col-0*. (**G**) *dgk4-1* PT showed reduced reproductive fitness when allowed to compete with the pollen of *Col-0* on emasculated *Col-0* flowers for fertilization. ‘*Col-0*’ and ‘*dgk4-1*’ represent seeds produced from self-fertilized *Col-0* and *dgk4-1* plants, and ‘*Col-0* + *dgk4-1*’ represents seeds produced from emasculated *Col-0* flower crossed with *Col-0* and *dgk4-1* pollen. The seeds from the cross (*n* > 70) screened on MS agar (1.1% w/v) containing 100 µg/mL of kanamycin, showed greater proportion (63.5%) of *Col-0* genotype.

In accordance with the in vitro germination phenotype, *in vivo* germinated PTs of *dgk4-1* are likewise growing consistently slower down the pistil than those of WT across all time points examined (Figures 1E and 1F). Importantly, the slowed PT growth of the mutant resulted in a significant reproductive fitness bias in favor of WT as observed in the crossing of emasculated WT flower with pollen from WT and *dgk4-1* that produced 63.5% of WT seeds when screened on MS agar containing 100 µg/mL kanamycin, the selective marker or the *dgk4-1* line used (Figure 1G), a result consistent with results reported for a different *dgk4* insertion line [11].

### DGK4 harbors a H-NOX-like center that yields NO-responsive spectral changes and catalytic activity

Through sequence analysis we have previously predicted that DGK4 contains a region spanning from H350 to R383 similar to heme centers of functional gas-responsive heme-NO/oxygen (H-NOX), heme-NO binding and NO-sensing families of proteins in other kingdoms [2]. In particular, this region harbors the HX[12]PX[14,16]YXSXR consensus pattern derived for heme *b* containing H-NOX centers in proteins from bacteria and animals and is present in plant orthologs in species such as poplar, castor bean and soybean but absent in other *Arabidopsis* DGKs (Figure 2A). The presence of the H-NOX-like signature suggests that DGK4 may accommodate a heme *b* and correspondingly, the diagnostic spectral properties should have a distinct response to NO. Recombinant DGK4 yields a Soret peak at 410 nm (Figure 2B), which is distinctly different from unbound hemin (protohemine IX: Soret band at 435 nm with a shoulder at 400 nm) and falls within the typical peak range observed for proteins with a histidine-ligated ferric heme *b* [13]. Reduction with sodium dithionite resulted in a red-shift of the Soret peak to 424 nm accompanied by the emergence of distinct *α* (558 nm) and *β* (526 nm) bands (Figures 2B and S3A). The ferrous state presumably represents the native state of DGK4 in the cytosol (*A. thaliana* cytosolic redox potential: −310 to −240 mV [14]). Exposure to air recovers the oxidized Soret peak (410 nm) of DGK4 after 20 min (Figure S3A). Importantly, addition of DEA NONOate attenuates the reduced Soret absorption (424 nm) in a concentration dependent manner hinting at the possibility of NO displacing the histidine ligand from the heme group (Figures 2C and S3B). This was suggested to be an essential step in the signaling of canonical H-NOX proteins [15]. Qualitatively, the observed spectroscopic behavior resembles that of canonical H-NOX proteins e.g., the H-NOX domain of *S. oneidensis* which showed Soret absorptions at 403 nm (ferric), 430 nm (ferrous) and 399 nm (ferrous, NO-bound) [16]. However, the frequencies and relative intensities of the observed Soret *α* and *β* peaks are indicative of a bis-histidine ligated heme *b* center as it is e.g. present in cytochromes *b5* or a heme-based cis-trans carotene isomerase Z-ISO [17]. In accordance DGK4 mutants which affect the heme binding site should result in a reduced heme absorption spectrum, a behavior which was e.g. demonstrated for Z-ISO [17]. According to this prediction, H350L and Y379L dgk4 mutants have reduced Soret band intensities of about 50% and 70% respectively (Figure 2D). Since the Soret bands were still present in the mutants albeit attenuated, we can expect a similar behavior in their reduction and NO spectra which we did observe with the H350L mutant protein (Figure S3). Overall, the H250L dgk4 mutant recorded a much larger decrease in reduced Soret bands than that observed with DGK4 WT at low NO donor concentration (0.25 mM DEA NONOate) while also requiring a slightly longer time (∼ 5 min more than DGK4 WT) to recover its oxidized Soret peak (410 nm) when exposed to air (Figure S3). Together with a marked reduced in Soret band intensity of DGK4 with point mutations at the H-NOX-like center, these results can be interpreted as the weakening of the heme environment.

**Figure 2.**
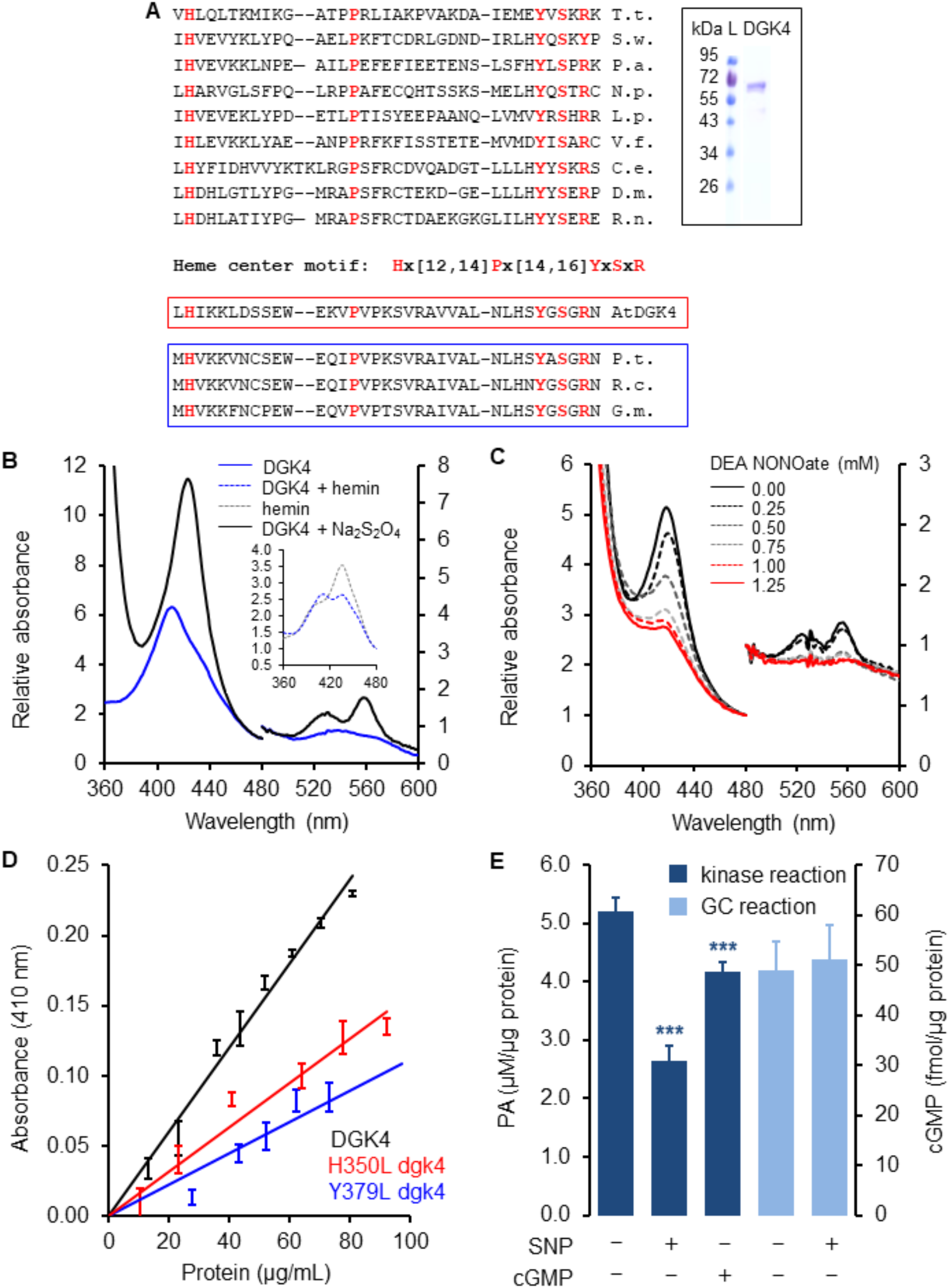
DGK4 harbors a H-NOX-like center and has NO-responsive spectral and catalytic activity. (**A**) The region H350 – R383 in DGK4 contains amino acid residues of annotated heme centers of gas-responsive proteins as shown in the alignment of *Thermoanaurobacter tengcongensis* (T.t.; GI: 3566245696), *Shewanella woodyi* (S.w.; Gl: 169812443), *Pseudoalteromonas atlantica* (P.a.; GI: 109700134), *Nostoc punctiforme* (N.p.; GI: 126031328), *Legionella pneumophila* (L.p.; GI: 52841290), *Vibrio fischeri* (V.f.; GI: 59713254), *Caenorhabditis elegans* (C.e.; GI: 52782806), *Drosophila melanogaster* (D.m.; GI: 861203), *Rattus norvegicus* (R.n.; GI:27127318), *Homo sapiens* (H.s.; GI: 2746083), *Arabidopsis thaliana* (A.t.; GI: 145359366), *Populus trichocarpa* (P.t.; GI: 224143809), *Ricinus communis* (R.c.; GI: 255581896) and *Glycine max* (G.m.; GI: 356567686) hemoproteins. The conserved functionally assigned residues of the heme centers are highlighted in red. Red box represents heme center of DGK4 and blue box represents orthologs of DGK4 that also contain similar heme centers. Inset: Recombinant DGK4 was generated and purified according to methods in Supplemental Experimental Procedures. (**B**) DGK4 (inset) contains a cytochrome *b*_*5*_ type heme center as indicated by the electronic absorption spectra in the ferric and ferrous state (Table S1). (**C**) UV-vis characterization of recombinant DGK4 reveals that NO attenuates the Soret peak of the ferrous heme center in a concentration dependent manner. (**D**) The H350L and Y379L dgk4 mutants have reduced heme binding. At 80 µg of protein, H350L and Y379L dgk4 have Soret peaks that are 0.5- and 0.3-fold of DGK4 (*n* = 3). (**E**), The kinase activity of DGK4 was reduced in the presence of SNP (1 mM) or cGMP (1 mM) but its GC activity was unaffected by SNP (1 mM). Kinase reaction mixture contains 40 mM Bis-Tris (pH 7.5), 5 mM MgCl_2_, 0.1 mM EDTA, 1 mM spermine, 0.5 mM dithiothreitol, 1 mM sodium deoxycholate, 0.02% (v/v) Triton X-100, 500 μM 1,2-DOG and 1 mM ATP and GC reaction mixture contains 50 mM Tris-HCl (pH 7.5), 1 mM GTP, 5 mM MgCl_2_ or MnCl_2_ (see Experimental Procedures for details). Dark blue bars represent kinase reactions while light blue bars represent GC reactions (*n* = 3). *\*\*\* = P* < 0.0005 compared to activity of DGK4 in the absence of SNP or cGMP.

Previously [2] we predicted DGK4 to be a bifunctional catalytic protein with (i) a canonical kinase domain capable of converting *sn*-1,2-diacylglycerol (DAG) with ATP into the corresponding phosphatidic acid (PA) and (ii) a moonlighting guanylyl cyclase (GC) activity that generates cGMP from GTP. Our prediction was recently confirmed by others [11]. We thus next focused on the catalytic activity of DGK4. NO and cGMP significantly inhibit DGK4 kinase activity but NO did not affect its GC activity (Figure 2E). Mutations in the H-NOX center did not affect the kinase activity of DGK4 as both H350L and Y379L *dgk4* mutants were functional and inhibited by NO to comparable degree as the WT (Figure S4). Having in mind the changes in the Soret band upon binding on NO in these mutated lines, this result could be interpreted as implying that NO signaling in PT chemotropic responses is achieved through alternative pathways. A diverse interpretation of this result could be ofered by considering that (i) while there is a reduction of intensity and slower recovery upon oxidation, there are still changes in the Soret band in the mutants, revealing some NO binding and (ii) enzymatic essays *in vitro*, with highly diluted enzyme concentrations and high concentrations of substrate hardly reproduce the cellular condition where molecular crowding determine specific kinetic properties [18], and the steady concentrations of NO may be much lower [8]. Our biochemical data also suggest that while the kinase activity of DGK4 seems to be inhibited by cGMP the GC activity of DGK4 is influenced by NO supporting an interpretation that the reduced NO response of *dgk4-1* PT is achieved primarily through a lipid/Ca^2+^ signaling pathway rather than through the activation of its GC moonlighting center as in soluble animal GCs. While consideration of a role for cyclic nucleotides in plant physiology is still affected by controversies regarding their synthesis and molecular targets, cGMP has been shown to activate Ca^2+^ conductivity by the CNGC18 channel localized in tip of PTs [19]. Taken together with the fact that DGK4 has been shown to localize in the cytosolic region of the PT tube apex [11], our results are consistent with a signaling role for DGK4 by transducing NO binding into lipid, cGMP, and Ca^2+^ pathways. Since DGKs catalyze the conversion of DAG to PA that is in turn essential for pollen growth [10] and in particular the mobilization of Ca^2+^ [7, 20], our results could be mechanistically interpreted first assuming that the response of DGK4 to NO signal alters its kinase activity resulting in lower cytosolic levels of PA and, reduced cytosolic Ca^2+^. These in turn are known to have various downstream signaling effects, namely in terms of ROP-GTPase, ion channel activation, resulting in vesicular trafficking and/or actin dynamics alterations and alteration of PT growth. In agreement with this interpretation, *dgk4-1* mutant PTs have recently been reported to exhibit altered mechanical properties with down-regulation of a cyclase associated protein, CAP1 which is involved in actin dynamics in addition to their reduced ability to target the ovaries *in vivo* [11]. Importantly, slow-down of PT growth has been deemed as optimizing the perception of chemical cues [21], and constitutes a plausible interpretation of the loss of chemotropic response in the mutant and likewise the seed-set reduction observed. As a diffusible gas, NO is well suited to perform fine-tuning of rapid cell-cell communications such as the pollen-stigma (2, 7). Here we propose DGK1 to be key in understanding the underlying signaling and NO-dependent cellular events during the progamic phase of sexual plant reproduction.

## Experimental Procedures

### Plant materials and growth conditions

Two mutant Arabidopsis lines (SALK_151239 and SALK_145081) with T-DNA insertions at the promoter of *DGK4* were purchased from Nottingham Arabidopsis Stock Center (NASC) and progenies were screened for homozygosity using PCR to detect mutant (T-DNA + *DGK4* reverse primer) and wild-type (*DGK4* promoter forward + *DGK4* reverse primer) chromosomes (Table S2). The homozygous mutant lines were subsequently referred to as *dgk4-1* and *dgk4-2* respectively. The T-DNA insertion sites were confirmed by sequencing (KAUST Bioscience Core Lab, Saudi Arabia). All seeds were cold stratified at 4 °C for 3 days. *Arabidopsis thaliana* (ecotype *Col-0*) and the T-DNA mutant lines were sowed on soil (Jiffy, USA) containing 50% (w/v) of vermiculite and grown in Percival growth chambers (CLF Plant Climatics, Germany) at 22 ± 2 °C and 60% of relative humidity under long day (16 hours light) photoperiod (100 *µ*M photons m^−2^ s^−1^).

### Characterization of *dgk4* mutant plants

RNA was extracted from pollen from approximately 300 flowers of WT and *dgk4-1* mutants (Qiagen, USA) and cDNA synthesized using SuperScript III reverse transcriptase according to manufacturer’s instructions (Invitrogen, UK). The cDNA was subjected to semi-quantitative RT-PCR with *DGK4* gene specific primers (Table S2) on an AB thermal cycler (Bio-Rad, USA) and *DGK4* gene expression was normalized against that of protein phosphatase 2A subunit A3, *PP2AA3* (At1g13320) (Table S2) using the ImageLab software (Bio-Rad, USA).

### *In vitro* pollen germination

*In vitro* pollen germination was performed as detailed previously [7, 8] in the absence or presence of NO provided by either SNP or DEA NONOate. In PT re-orientation experiments, pollen was allowed to germinate for 2 hours before image acquisition at specified intervals. The growth of at least 100 PTs were measured on a Nikon Eclipse TE2000-S inverted microscope equipped with an Andor iXon3 camera across a range of pH (pH 6.5 to 8.5) and in optimal (5 mM) and low (100 µM) Ca^2+^ media. In NO-treated pollen germination and PT growth experiments, at least 150 unique pollen/PTs were considered. Image frames covering the entire growth area of the culture dish that is mounted on automated stage, were acquired using the Nikon Eclipse TE2000-S inverted microscope which is equipped with a Hamamatsu Flash28s CMOS camera. The PT lengths were measured using NeuronJ [22]. In PT re-orientation studies, PTs were challenged with a NO probe provided by a glass pipette tip with a ∼ 5 µm aperture that was pre-filled with 10 mM SNP-agarose (1%) and placed 60 µm away from the growing PT tip using micromanipulators with a stepper-motor-driven three-dimensional positioner. The growth and bending response of four randomly selected healthy growing PTs were monitored by real-time imaging using a Nikon Eclipse TE300 inverted microscope equipped with an Andor iXon3 camera and the bending angles measured using ImageJ [23].

### Reproductive fitness test

The reproductive fitness of *dgk4-1* was examined by crossing emasculated WT (ecotype *Col-0*) flowers with pollen from both WT and *dgk4-1*. The desiccated seeds were collected, surface sterilized and stored at 4 °C for 3 days before growing on MS agar (1.1% w/v) containing 100 µg/mL kanamycin (Sigma-Aldrich, St. Louis, MO). The proportion of WT to *dgk4-1* seeds were scored after 7 days of growth.

### UV-visible absorption spectroscopy

The UV-visible spectra of affinity purified recombinant DGK4 (200 µg/mL) was recorded on a PHERAstar FS micro-plate reader (BMG Labtech, USA). The heme environment of DGK4 was characterized by the addition of a reducing agent, sodium dithionite (Na_2_S_2_O_4_) to a final concentration of 10 mM and absorbance was immediately measured and examined for spectral changes. The protein sample was then exposed to air and any recovery of the oxidized peak was monitored by the same spectra measurements at 5 min intervals. The heme-NO complex was generated by immediately adding the NO donor DEA NONOate to a pre-reduced recombinant DGK4 before making the same spectral measurements.

### DAG kinase and GC assays

DAG kinase assay and phospholipid extraction was performed using 30 μg purified recombinant protein in a reaction mixture containing 40 mM Bis-Tris (pH 7.5), 5 mM MgCl_2_, 0.1 mM EDTA, 1 mM spermine, 0.5 mM dithiothreitol, 1 mM sodium deoxycholate, 0.02% (v/v) Triton X-100, 500 μM 1,2-DOG and 1 mM ATP, in the absence or presence of 1 mM SNP or 0.65 mM DEA NONOate. PA generated from the reactions was measured using the Total Phosphatidic Acid Assay kit according to the manufacturer’s instructions (Cayman Chemical, Michigan USA).

GC assay was performed using 10 µg purified recombinant protein in a reaction mixture containing 50 mM Tris-HCl (pH 7.5), 1 mM GTP, 5 mM MgCl_2_ or MnCl_2_, in the absence or presence of 1 mM SNP or 0.65 mM DEA NONOate. cGMP generated from the reactions was measured using the cGMP enzyme immunoassay (EIA) Biotrak System with the acetylation protocol according to the manufacturer’s instructions (GE Healthcare, Illinois USA).

### Chemicals and statistical analysis

All chemicals were purchased from Sigma unless stated otherwise. Statistical analysis was performed using Student’s *t*-test with Microsoft Excel 2010. Significance was set to a threshold of *P* < 0.05 and *n* values represent number of biological replicates.

PT growth *in planta* and protein expression and purification are described in detail in Supplemental Experimental Procedures.

## Supplemental Information

Supplemental Information contains four figures, two tables, supplemental experimental procedures and supplemental references.

## Author Contributions

C.G. conceived of the project and A.W., L.D. and M.T.P. conducted the experiments. All authors contributed to the data analyses and writing of the manuscript.

## Acknowledgments

This research was supported by the King Abdullah University of Science and Technology (CG lab) and National Science Foundation grants MCB-1616437 and 1714993 (JF lab). A.W. is supported by the National Natural Science Foundation of China (Grant no. 31850410470) and the Zhejiang Provincial Natural Science Foundation of China (Grant no. LQ19C130001).

