## Supplemental materials for "The *Arabidopsis* Diacylglycerol Kinase 4 is involved in nitric oxide-dependent pollen tube guidance and fertilization"

### Supplemental Information

**Figure S1**

Characterization of *dgk4* T-DNA insertion mutant plants

**Figure S2**

Homozygous *dgk4-2* PT has slower growth rate and reduced NO-dependent growth response

**Figure S3**

DGK4 harboring point mutation at the H-NOX-like center yields spectral behavior similar to WT

**Figure S4**

Kinase activities of DGK4 and mutant dgk4 harboring point mutations at the H-NOX-like center were inhibited by NO

**Table S1**

UV-Vis spectroscopic data of selected heme proteins

**Table S2**

Primers for cloning of *DGK4* and characterization of *dgk4-1* and *dgk4-2* plants

**Movie S1**

PT re-orientation responses of *Col-0* and *dgk4-1* to NO

### Supplemental Experimental Procedures

### Supplemental References

### Figure S1. Characterization of *dgk4* T-DNA insertion mutant plants

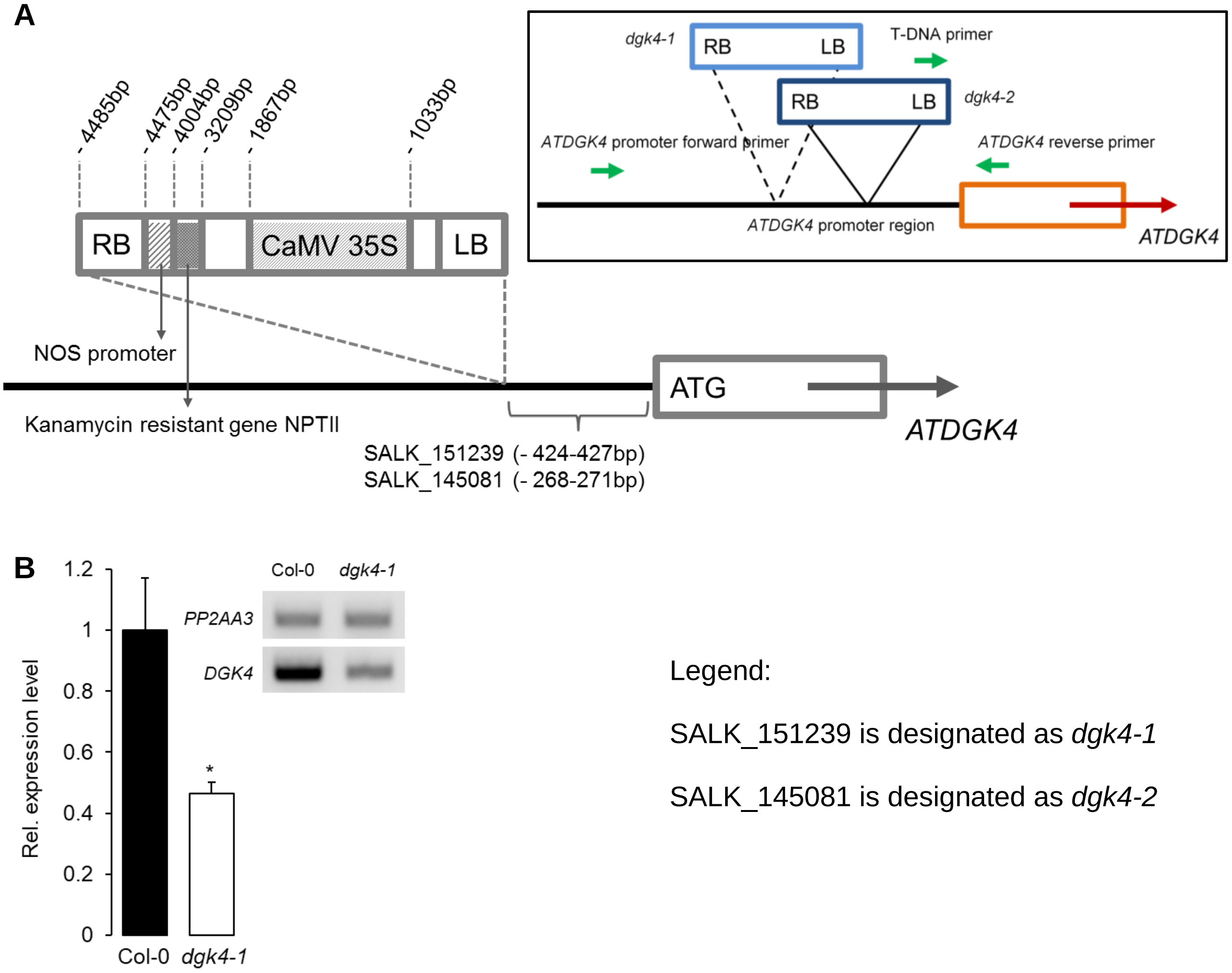


(**A**) Schematic view of T-DNA insertion sites of *dgk4* mutants. Location and content of SALK T-DNA insertions are labelled. LB and RB indicate Left Border and Right Border of the T-DNA respectively. Inset: Green arrows indicate the position and direction of primers (see also Table S2) used in RT-PCR to determine *DGK4* expression levels. (**B**) *dgk4-1* pollen has reduced *DGK4* mRNA levels as estimated by semi-quantitative RT-PCR. * = $P < 0.05$ compared to *DGK4* mRNA levels of *Col-0* pollen and gel pictures are representative of three independently derived biological replicates.

**Figure S2. Homozygous *dgk4-2* PT has slower growth rate and reduced NO-dependent growth response**

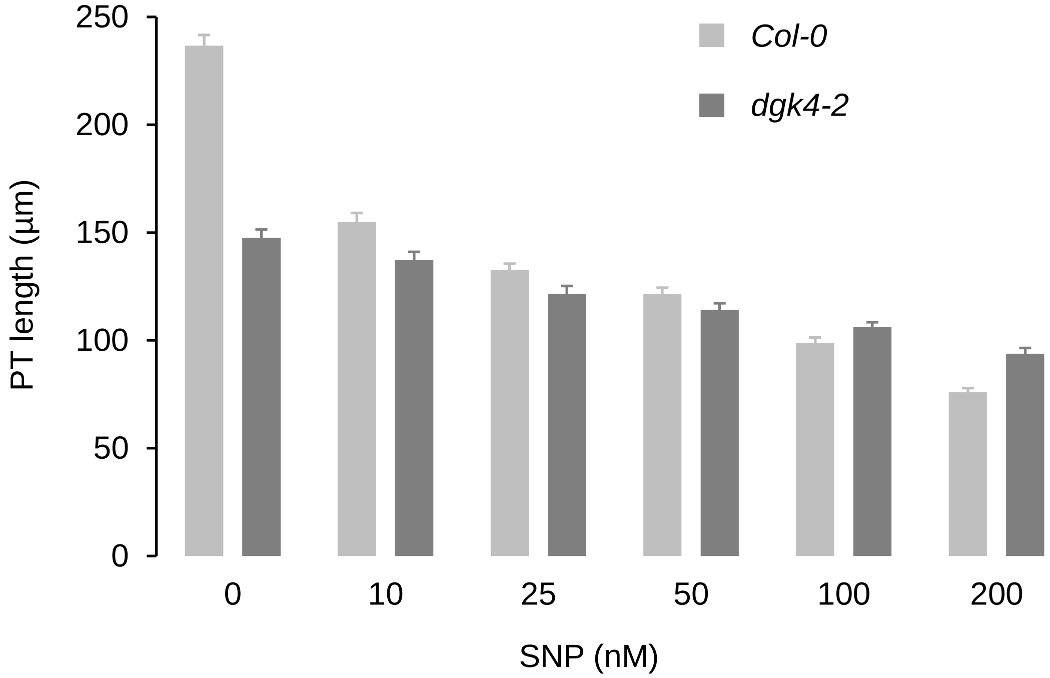


NO-dependent inhibition of *dgk4-2* PT growth is reduced compared to that of *Col-0*. NO was provided by either SNP. *In vitro* pollen germination was performed as detailed previously [1, 2] and PT length was analyzed by capturing images covering the entire growth area of the culture dish that is mounted on an automated stage using the Nikon Eclipse TE2000-S inverted microscope equipped with a Hamamatsu Flash28s CMOS camera. The pollen tube lengths were then measured using NeuronJ [3]. Error bars represent standard error of the mean ($n$ > 150).

**Figure S3. DGK4 harboring point mutation at the H-NOX-like center yields spectral behavior similar to WT**

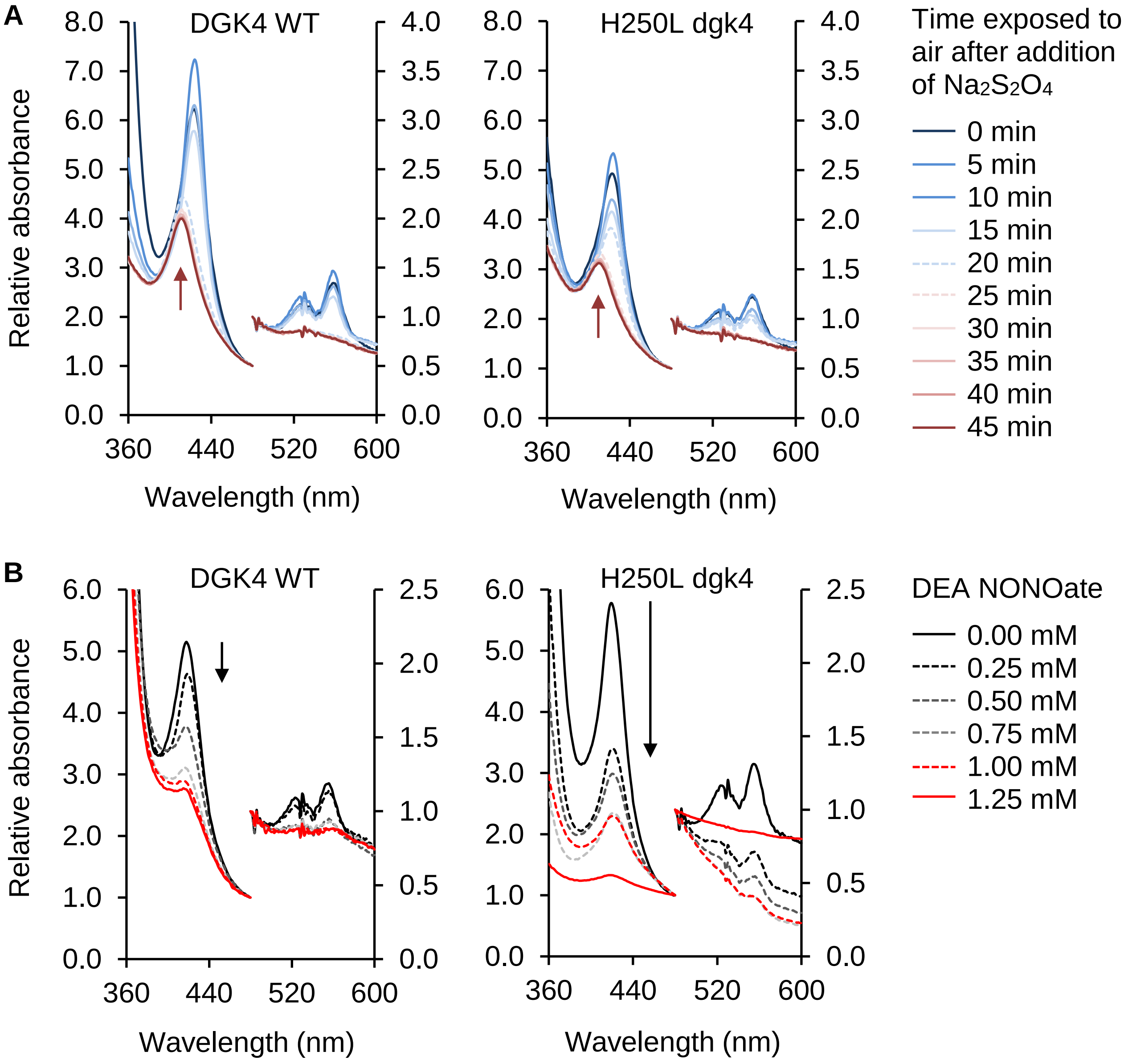


(**A**) UV-vis characterization reveals that the Soret peaks (410 nm) of 80 µg DGK4 WT and H250L dgk4 mutant proteins were both red-shifted to 424 nm accompanied by the emergence of distinct $\alpha$ (558 nm) and $\beta$ (526 nm) bands when reduced with sodium dithionite. The oxidized Soret peaks (410 nm) (red arrows) of both DGK4 WT and H250L dgk4 mutant were fully recovered after 20 and 25 min of exposure to air respectively. (**B**) Addition of DEA NONOate to reduced DGK4 and H250L dgk4 attenuates the Soret absorption (424 nm) in a concentration dependent manner where the Soret, *β*- and *α*-peaks vanish with increasing concentration of the NO donor. H250L dgk4 mutant recorded a much larger decrease in reduced Soret bands than that observed with DGK4 WT at low NO donor concentration (0.25 mM DEA NONOate) (black arrows) while also requiring a slightly longer time (~ 5 min more than DGK4 WT) (red arrows) to recover its oxidized Soret peak (410 nm) when exposed to air.

**Figure S4. Kinase activities of DGK4 and mutant dgk4 harboring point mutations at the H-NOX-like center were inhibited by NO**

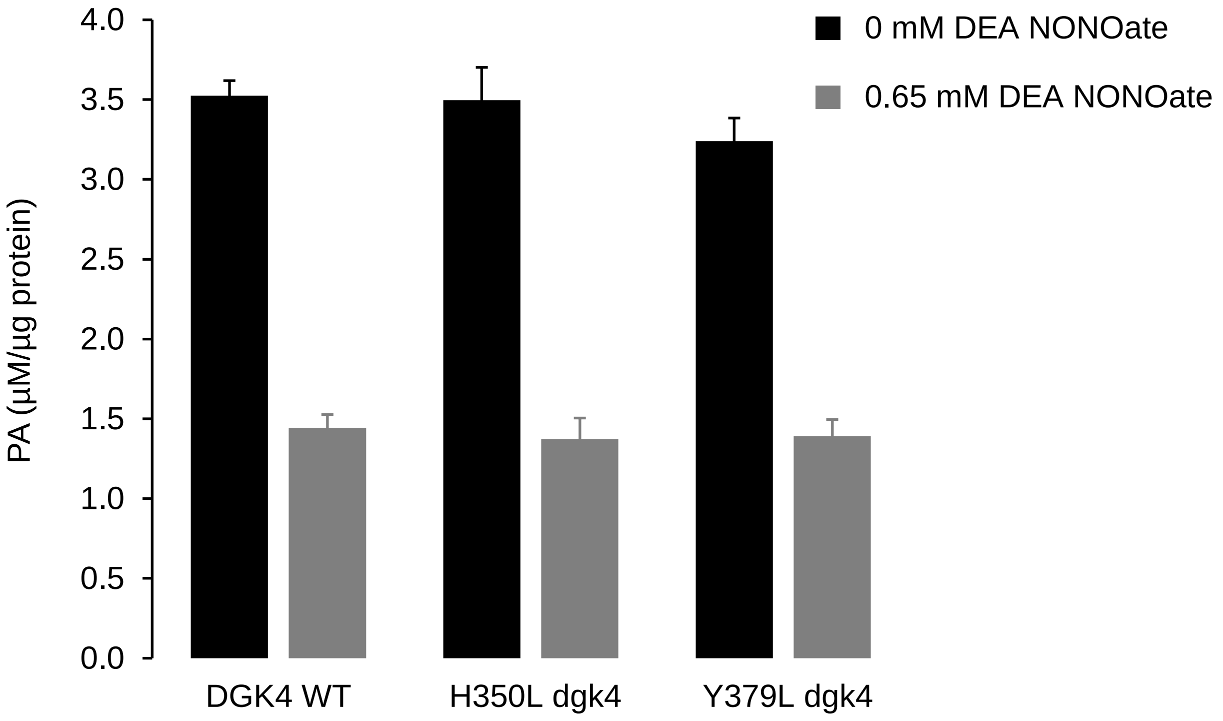


The kinase activities of dgk4 mutants H350L and Y379L were unaffected by the point mutations at the H-NOX-like center and were inhibited by NO to comparable degree as the WT. Kinase assay was done in reaction mixtures containing 40 mM Bis-Tris (pH 7.5), 5 mM $MgCl_2$, 0.1 mM EDTA, 1 mM spermine, 0.5 mM dithiothreitol, 1 mM sodium deoxycholate, 0.02% (v/v) Triton X-100, 500 µM 1,2-DOG and 1 mM ATP, with or without 0.65 mM DEA NONOate (see Experimental Procedures for details). Black solid bars represent kinase reactions performed in the absence of NO while grey solid bars represent kinase reactions performed in the presence of NO ($n$ = 6).

**Table S1. UV-Vis spectroscopic data of selected heme proteins**

| | Soret / $\beta$ / $\alpha$ ; max. absorption / nm | | | Source |
|---|---|---|---|---|
| Protein (*sp.*) | ferric | ferrous | ferrous-NO | |
| DGK4 (*At*) | 410 / $\alpha + \beta$ 534 | 424 / 526 / 558 | 418 / - / - | This work |
| H-NOX (*So*) | 403 / - / - | 430 / $\alpha + \beta$ 560 | 399 / 543 / 572 | [4] |
| Cyt b5 (*Gl*) | 411 / $\alpha + \beta$ 532 | 423 / 526 / 558 | | [5] |
| Z-ISO (*At*) | 414 / $\alpha + \beta$ 531 | 414 / 529 / 559 | | [6] |

Legend: *At*: *Arabidopsis thaliana, So: Shiwanella oneidensis, Gl: Giardia lamblia.*

**Table S2. Primers for cloning of *DGK4* and characterization of *dgk4-1* and *dgk4-2* plants**

| Primer name | Sequence (5' – 3') |
|---|---|
| ***DGK4* cloning** | |
| *DGK4* F | ATGGAATCACCGTCGATTGG |
| *DGK4* R | TCAATCTCCTTTGACGACCAA |
| *DGK4* H-L F | TTATGACATTGCTTATAAAAAAGTTGG |
| *DGK4* H-L R | CAACTTTTTTATAAGCAATGTCATAAA |
| *DGK4* Y-L F | ATCTACATAGCTTAGGAAGTGGAAGAA |
| *DGK4* Y-L R | TCTTCCACTTCCTAAGCTATGTAGATT |
| **Screening for homozygous *dgk4-1* and *dgk4-2* plants** | |
| *DGK4* promoter forward | TGTTTCTGACATCTGAGAACTTTT |
| *DGK4* reverse | GATTGCATTCTTCGTAAAGACG |
| *T-DNA* | GTTCACGTAGTGGGCCATCG |
| **Expression of *DGK4*** | |
| *DGK4* qPCR forward | CGTCGATTGGTGATTCATTG |
| *DGK4* qPCR reverse | TTGCAATGCGGAGATATTGA |
| *PP2AA3* qPCR forward | GCGGTTGTGGAGAACATGATACG |
| *PP2AA3* qPCR reverse | GAACCAAACACAATTCGTTGCTG |

Note: The underlined nucleotides incorporate the mutations changing histidine or tyrosine residues at positions 350 and 379 to leucine.

**Movie S1. PT re-orientation responses of *Col-0* and *dgk4-1* to NO**

Movie file attached separately.

### Supplemental Experimental Procedures

#### PT growth *in planta*

PT growth in the pistil of hand pollinated WT and *dgk4-1* plants was examined by collecting the pistils at different time points (3 – 8 hours) after pollination. Aniline blue staining of PTs in the pistil was performed as described previously [7] and PT length in the pistil was measured using ImageJ [8].

#### Protein expression and purification

A Gateway compatible clone (DKLAT5G57690.1) containing the full-length coding sequence of *DGK4* was purchased from Arabidopsis Biological Resource Center (ABRC). The DGK4 sequence was recombined into the pDEST17 His-tagged expression vector and transformed into *E. coli BL21 A1* (Invitrogen, USA). Expression of recombinant DGK4 was induced with 0.2% (w/v) L-arabinose. Cells were lysed in a guanidium lysis buffer and the supernatant loaded onto a Ni-NTA agarose column for affinity purification under denaturing conditions using urea-containing buffers. Denatured recombinant DGK4 was re-folded by gradual dilution of urea in a linear gradient using an AKTA FPLC (GE Healthcare, UK). Hemin (30 µg/mL) was added to the re-folding buffers to allow for incorporation of heme into DGK4 as it assumes native conformation. Excess hemin was removed by size exclusion and recombinant DGK4 stored in ‘Buffer’ containing 20 mM $Na_2H_2PO_4$, 500 mM NaCl, 500 mM sucrose, 100 mM non-detergent sulfobetaines (NDSB), 0.05% (w/v) polyethylene glycol (PEG), 4 mM reduced glutathione, 0.04 mM oxidized glutathione and SIGMAFAST protease inhibitor cocktail (1 tablet per 100 mL solution).

Two single dgk4 mutants (H350L and Y379L) were constructed using site directed mutagenesis by PCR [9]. To construct the H350L dgk4 mutant, two overlapping fragments of the DGK4 coding sequence both incorporating the mutation, were amplified from the pDEST17-*DGK4* plasmid using the respective *DGK4* F and *DGK4* H-L R (for 1$^{st}$ fragment amplification), and *DGK4* H-L F and *DGK4* R (for 2$^{nd}$ fragment amplification) primer pairs (Table S2). The two overlapping fragments both incorporating the mutations were then used as templates for a PCR reaction using the full-length *DGK4* F and *DGK4* R primer pairs (Table S2) which generated a full-length dgk4 H350L mutant sequence. The dgk4 Y379L mutant was generated using the same method but with the following mutagenic primers pairs, *DGK4* F and *DGK4* Y-L R, and *DGK4* Y-L F and *DGK4* R (Table S2). The *DGK4* mutant PCR products were inserted into the PCR8/GW/TOPO vector (Invitrogen, USA) by TA cloning, recombined into the pDEST17 His-tagged expression vector and transformed into *E. coli BL21 A1* (Invitrogen, USA). Mutant dgk4 was expressed and affinity purified in the same manner as DGK4.
